# Visual space curves before eye movements

**DOI:** 10.1101/2021.10.13.464140

**Authors:** Ifedayo-EmmanuEL Adeyefa-Olasupo, Zixuan Xiao

## Abstract

In most experiments within the field of cognitive and systems neuroscience, fixation is often required prior to the onset of an experimental trial. However, the term “fixation” is rather misleading since our eyes are constantly moving. One type of transient miniature movement ubiquitously observed during fixation is commonly referred to as fixational eye movements or microsaccades. Perimicrosaccadic compression of visual space — the ability of retinotopic cells to transiently exhibit predictive spatiotemporal retinotopic compressive shifts toward the target of an impending microsaccade — is known to dramatically alter visual perception. However, whether perimicrosaccadic compressive shifts can become spatially asymmetric, that is, continuously directed toward a specific foveal region over another (e.g., an upper over a lower region in the fovea) and for what purpose, remains poorly understood. Assuming that these transient shifts are indeed asymmetric under certain conditions, the perceptual and oculomotor consequences such asymmetricity might accompany across visual space is poorly understood. Equally unaccounted for is a mechanistic account of the neural computation and architecture that could support these transient asymmetric shifts while the visual system actively maintains retinotopic organization. Here, we systematically measured visual sensitivity in human subjects to faint probes presented during fixation and around the time of saccadic eye movement at geometrically symmetric retinotopic locations in the foveal, parafoveal, and peripheral regions of visual space. Remarkably, we observed transient local asymmetric visual sensitivity differences between these symmetric retinotopic locations where none should be observed. Equally surprising, we observed the trajectories of saccadic eye movements, which are expected to travel along a linear path, routinely deviate along a curved path toward orthogonal eccentric locations. To provide a mechanistic account of the neural computation and architecture that may explain our results, we proposed a novel neurobiologically inspired phenomenological force field model in which underlying attentional and oculomotor signals are modulated by transient eccentric error signals that manifest as temporally overlapping predictive forces and impinge on the retinotopic visual cortex. These forces, which transiently bias putative population sensitivity toward an orthogonal retinotopic foveal region and, around the time of a saccadic eye movement, along an axis orthogonal to the saccade direction toward a mislocalized peripheral region, succinctly capture the essence of our empirical observations.

## INTRODUCTION

The central 2° of the visual space is extensively represented in the retina, the superior colliculus (SC), and the primary visual cortex (V1) and is thus well suited for tasks that necessitate a detailed exploration of everyday visual objects (e.g., fruits). In the event that a visual object of interest lands on the peripheral retina, a large and rapid eye movement, known as a saccade, is recruited by the visual system to bring that object into the central 2° of visual space (i.e., to foveate). To explore the fine visual details that might convey the state of the foveated object (e.g., its ripeness or aposematic extent), microsaccades, also referred to as small fixational eye movements, are continuously being executed by the visual system while the fine visual features of the object are brought in and out of the foveola, the most central 1° of visual space where visual resolution is the highest (*1, 2, 3*).

Prior to the onset of a fixational eye movement, marked by a local transient shift in gaze, retinotopic cells in the superior colliculus predictively shift their neural resources toward the target of an impending fixational eye movement (*4*). This phenomenon is known as perimicrosaccadic compression of visual space. On the perceptual domain, perimicrosaccadic compression, which enhances foveal representation, causes human subjects to erroneously mislocalize eccentric faint stimuli as more foveal (*5,6*). Remarkably, perimicrosaccadic compression is mechanistically consistent with perisaccadic compression of visual space – the ability of retinotopic cells to transiently exhibit predictive spatiotemporal retinotopic compressive shifts toward the target of the saccade (*7*). On the perceptual domain, these transient retinotopic convergent shifts cause human subjects to also mislocalize flashed faint probes; however, in this case, toward the target of the saccade (*8*). Previous studies have shown that perisaccadic compressive shifts are not always symmetric (*9, 10*). This leads to frequent orthogonal mislocalization of flashed faint stimuli away from the saccade target (*11, 12*). However, in the case of perimicrosaccadic compression, it is unknown whether these compressive shifts can also demonstrate asymmetric biases (e.g., persistent and sustained compressive retinotopic shifts toward a selected foveal region over another region, which constitutes the destination of successive fixational eye movements). If perimicrosaccadic compression can become asymmetric, on the perceptual domain, this should result in transient increases in visual sensitivity at retinotopic foveal locations that either overlap with or are proximal to the foveal region that constitutes the target of these asymmetric perimicrosaccadic compressive shifts (*13*). Furthermore, assuming this perceptual bias predicts what we might expect in the fovea, performance in the parafoveal and peripheral regions of visual space is harder to predict and unsurprising remains unexplored. In addition, no study to date has been able to provide a mechanistic account of the neural computation and architecture the visual system uses to mediate asymmetric perimicrosaccadic compression of visual space while controlling for any overcompression toward the fovea (i.e., maintaining retinotopic organization).

Equally unaccounted for are two issues pertaining to the perceptual and motor influences perimicrosaccadic compression in general wield around the time of a saccade. Perimicrosaccadic shifts precede perisaccadic compressive shifts. However, it is poorly understood the extent to which these two forms of compression temporally overlap or take place within distinct temporal windows, along with the perceptual effect that such temporal dynamics accompany around the time of the saccade. On the motoric domain, studies have shown that saccades that are expected to travel along a linear path can deviate along a curved trajectory (*14*). While perimicrosaccadic compression causes subjects to mislocalize faint visual stimuli, which temporally overlap with the planning and preparation of a saccade, it is also unknown whether the spatial extent of these compressive shifts can also cause subjects to mislocalize the saccade target, which, if it is the case, should fundamentally influence the planning and trajectory of the saccade (*15*).

To provide an account for (i) the spatiotemporal perceptual consequences across visual space that are driven by transient asymmetric perimicrosaccadic compressive shifts, (ii) how the planning and trajectory of a saccade are influenced by this asymmetricity, and (iii) whether this asymmetricity influences perisaccadic compressive shifts and the perceptual signatures they accompany, we first assessed saccade landing positions after subjects executed a saccade. This informed us as to where subjects perceived the fixed saccade target to be located during fixation. Following this, we assessed the transient consequences of visual sensitivity to visual probes that were presented around the path of a saccade in the foveal, parafoveal, and peripheral retinotopic regions of visual space during fixation and around the time of a saccade. In addition, saccade trajectories were investigated to assess whether the planning of a saccade and its eventual execution along with observed transient changes in visual sensitivity share an underlying cause. Next, we investigated the rhythmicity that actively supported these transient changes in visual sensitivity to assess whether they resemble known rhythmic attentional signatures that mediate neural activity within retinotopic brain areas. Finally, we proposed a novel neurobiologically inspired phenomenological force field model that succinctly recapitulated transient changes in visual sensitivity at less (foveal) and more eccentric (parafoveal) retinotopic regions of visual space.

In a dimly lit room, during fixation, flashed low-contrast probes (at a contrast level that the subject could detect with 50% accuracy) were randomly flashed for 20 ms at 2.5 degrees of visual angle from the fixation dot (*Experiment 1*), from the midpoint (at an eccentricity of 5°) between the fixation dot and the fixed saccade target (*Experiment 2*), or from the fixed saccade target (*Experiment 3*), at tangential retinotopic locations (i.e., retinotopic locations around the presumed path of a saccade, Fig. 1A), geometrically symmetric to one another with respect to the radial axis (i.e., the presumed path along which subjects should plan their impending saccade). Temporally, low-contrast probes were flashed either during fixation or after the extinguishment of the fixation dot, which served as the central motor cue for subjects to make a planned saccade toward the retinotopic location along which they perceived the fixed saccade target to be positioned (see Methods, Fig. 1B). With respect to target onset, flashed probes fell within the time window of 0 to 1285 ms after target onset. The task of each subject was to report with a button press whether they were able to detect a flashed probe while making a saccade toward the fixed saccade target. Note that unlike previous psychophysical studies that only measured a single tangential axis, aside from the fixed saccade (or eccentric) target, we did not present any secondary salient spatial cue along a preselected tangential axis (*16, 17*). This ensured that if any differences between these tangential axes were to be observed, they could not be attributed to an experimental artifact. In addition, to control for any strategic restriction of attentional and oculomotor resources along a single tangential axis, probes were isotropically flashed along the radial axis. Finally, to control for any false responses, in 25% of all experimental trials, no probes were flashed, which yielded a false alarm rate of 1.2% along the tangential axes. Indeed, because of the aim of our study highlighted above, perceptual and oculomotor responses obtained in both control and radial conditions were discarded and are therefore not included in the main analyses reported below.

**Fig. 1.**
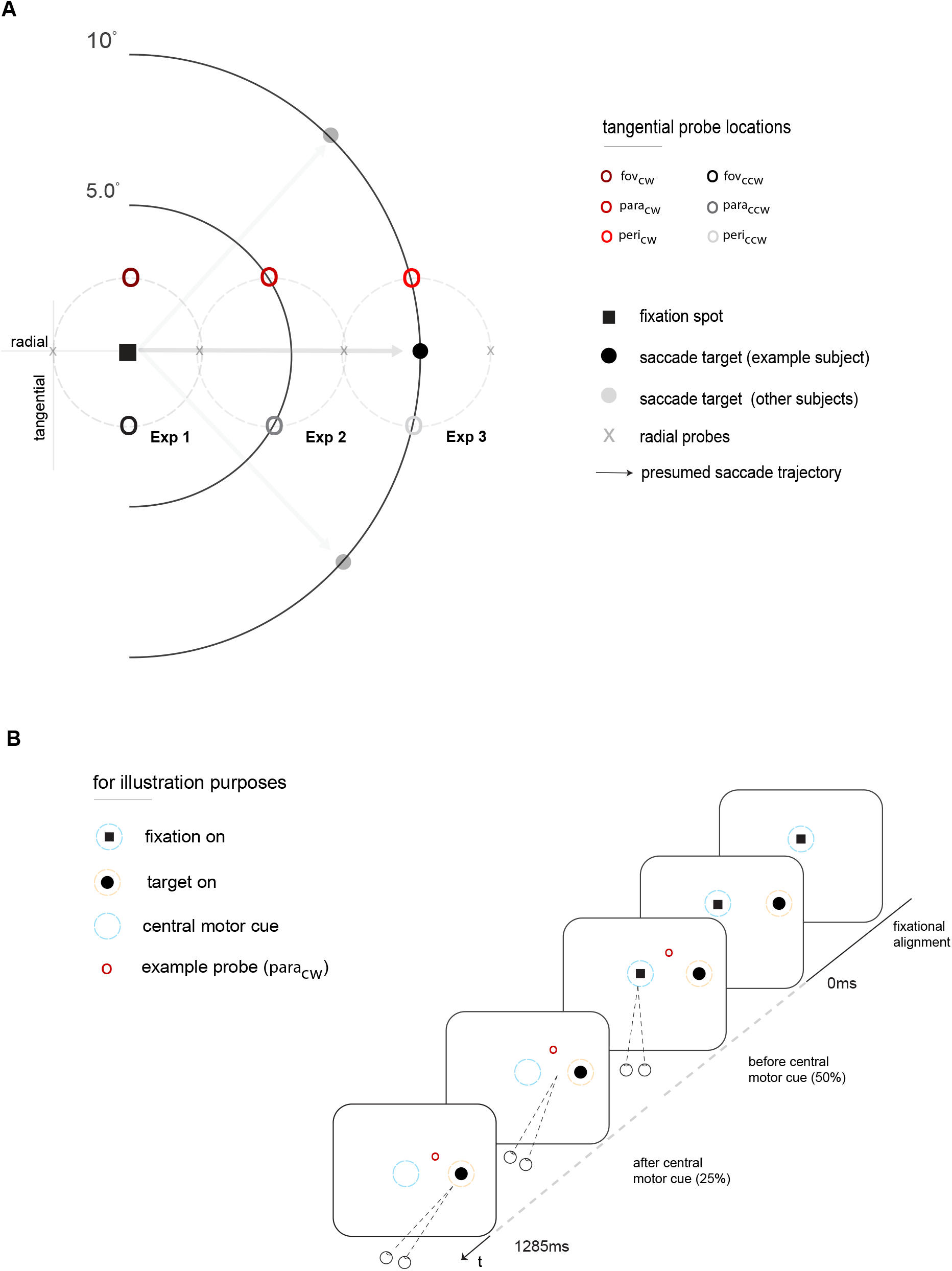
Spatiotemporal profile of experiments. **(A)** Markers in the illustration of visual space show all retinotopic locations at which a probe was flashed before and after the deployment of attention, saccade planning, and, finally, the execution of a saccade. Probes of interest, presented along the tangential axes, are conspicuously highlighted in open circle markers. **(B)** Temporal sequence of a valid trial across experiments with respect to target onset.

## RESULTS

### Local transient shifts in gaze and saccade landing positions

Across experiments, after the onset of the fixed eccentric target, we observed local transient shifts in gaze (*1*). However, specifically from 0 to 209 ms, these local transient shifts in gaze were unexpectedly averted away from the retinotopic location of the fixed eccentric target and were uniquely biased toward a foveal region that is more proximal to the foveal tangential clockwise *fov*_*cw*_ location (Fig. 2A). This observation aligns with previous studies that report that some attentional resources, but not all, are allocated away from the preselected eccentric spatial cue shortly after it appears on the peripheral retina (*18, 19*). From 210 ms to 820 ms after the onset of the fixed eccentric target (Fig. 2B), we observed that these transient shifts in gaze continued to remain eccentric but were now being compressed in a persistent and sustained manner toward a foveal region more proximal to the foveal tangential counterclockwise *fov*_*ccw*_ location. These persistent and sustained compressive shifts in gaze are in agreement with a seminal study that reported enhanced visual processing at the attended foveal site (*20*), thereby causing visual receptive fields to compress around this region in the fovea. The spatial distribution between these local transient shifts in gaze along the radial and tangential axes was statistically significant (*rad*_*shift*_, p = 4.0058e-36; *tan*_*shift*_, p = 0.0149).

**Fig. 2.**
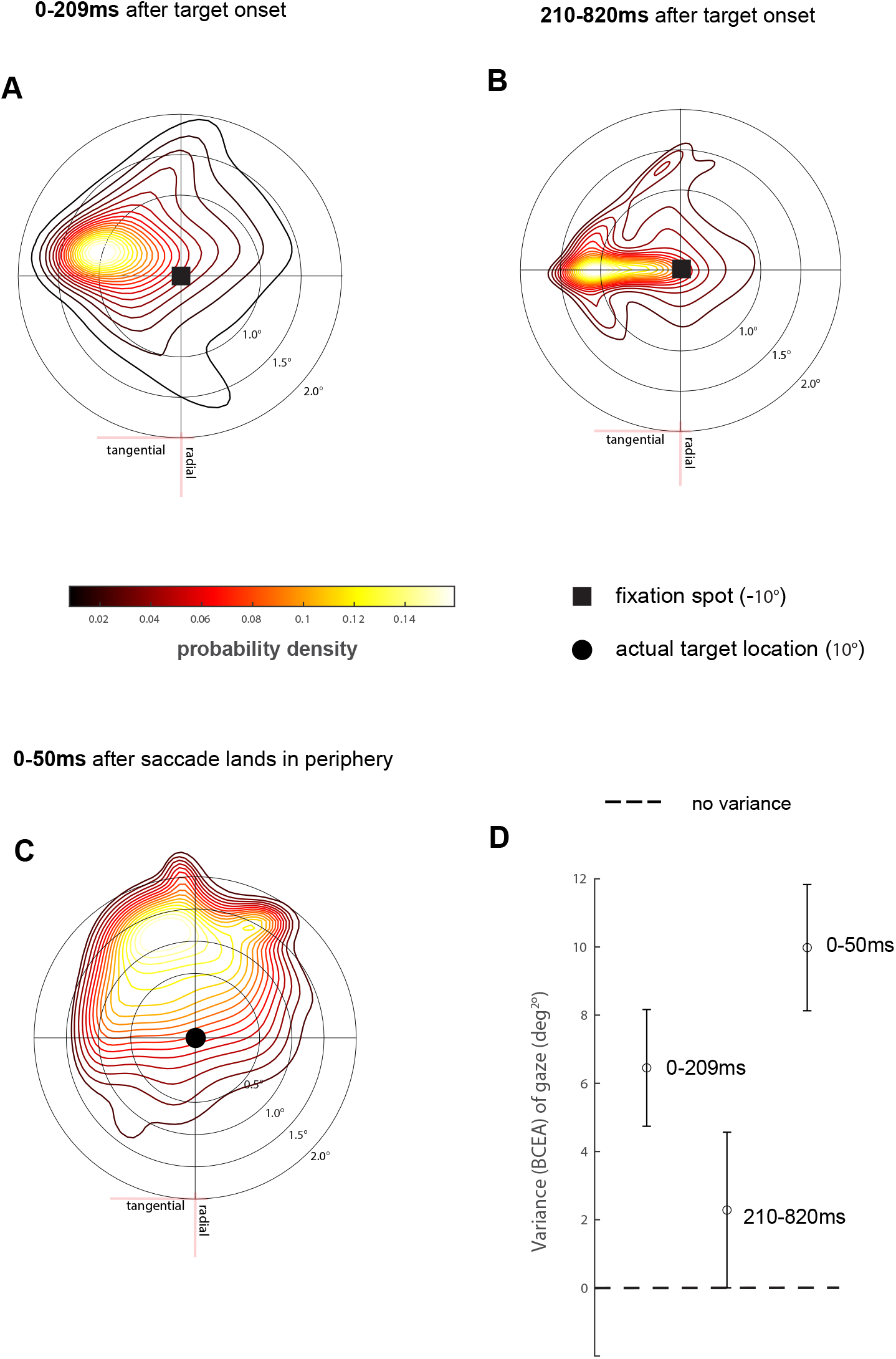
Gaze dynamics, saccade landing position, variance estimates. **(A-D)** Probability density map of local shifts in central gaze and saccade landing position was calculated using a bivariate gaussian kernel along with their bivariate contour ellipse area estimates for each probability density map. Error estimates are standard error of means across observers.

Across experiments, we observed that saccade landing positions, from 0 to 50 ms after saccade offset (i.e., the moment the eye lands in the periphery), were dramatically mislocalized toward the peripheral tangential clockwise *peri*_*cw*_ location (Fig. 2C). Because the visual system, as it plans an impending saccade, must estimate the spatial extent of the eccentric saccade target that is concurrently being sampled on the peripheral retina during fixation (*15, 21, 22*), it is highly likely that subjects erroneously perceived the fixed eccentric saccade target in the direction of the *peri*_*cw*_ location just before saccade onset (*11, 12*). As expected, a larger variance was observed for the initial locus of gaze (6.84 deg^2^) than for the variance associated with later compressed shifts in gaze (2.12 deg^2^). For the variance associated with the saccade landing positions, a larger variance (10 deg^2^) was observed when compared to the variance obtained for the initial and later shifts in gaze (Fig. 2D).

### Asymmetric foveal compression

As our eyes incessantly move during fixation, visuomotor signals arrive faster at the target of impending fixational eye movement in the direction of the more eccentric retinotopic regions. Consequently, retinotopic foveal locations that are more proximal to the target of impending fixational eye movement inherently benefit from earlier and more enhanced visual processing (20, *23, 24*). Conversely, in the more eccentric regions of visual space, visual processing is significantly delayed, which results in an erroneous representation at more eccentric locations (*9, 10, 11, 12, 25*). With this in mind, an account emerges that may explain why subjects frequently mislocalized the fixed eccentric target toward the tangential clockwise axis (Fig. 2C). Specifically, we posit that just after the onset of an eccentric target, which is accompanied by initial inward radial-tangential shifts in gaze (Fig 2A), some amount of visual resources at the more eccentric retinotopic region of visual space is allocated toward the target of a fixational eye movement (*18, 19, 20, 23, 24*). Consistent with the prediction of perimicrosaccadic compression of visual space, an eccentric flashed stimulus within this presaccadic window (>100 ms to saccade onset) would most likely appear more foveal (Fig. 3A, *5*). Furthermore, we posit that accompanying these transient inward radial-tangential shifts are transient asymmetric compressive components that are accompanied here by persistent and sustained compressive shifts in gaze toward the tangential counterclockwise axis (Fig. 2B). These sustained transient compressive shifts require resources already present in the fovea (at the *fov*_*cw*_ location) and additional resources from the more eccentric regions of visual space (*20*). Because of this demand, the more eccentric regions of visual space, keeping only residual resources centripetally distributed around a mislocalized preselected eccentric target (*9, 11*), are uniquely devoid of the visual resources they need to calibrate their representations initially distorted by transient inward radial-tangential compressive shifts. Consequently, within a presaccadic window, this results in an erroneous (i.e., curved) representation of visual space, marked by an orthogonally biased gaze that is radially connected to an orthogonally mislocalized target (Fig 3B, *26, 9, 10*).

**Fig. 3.**
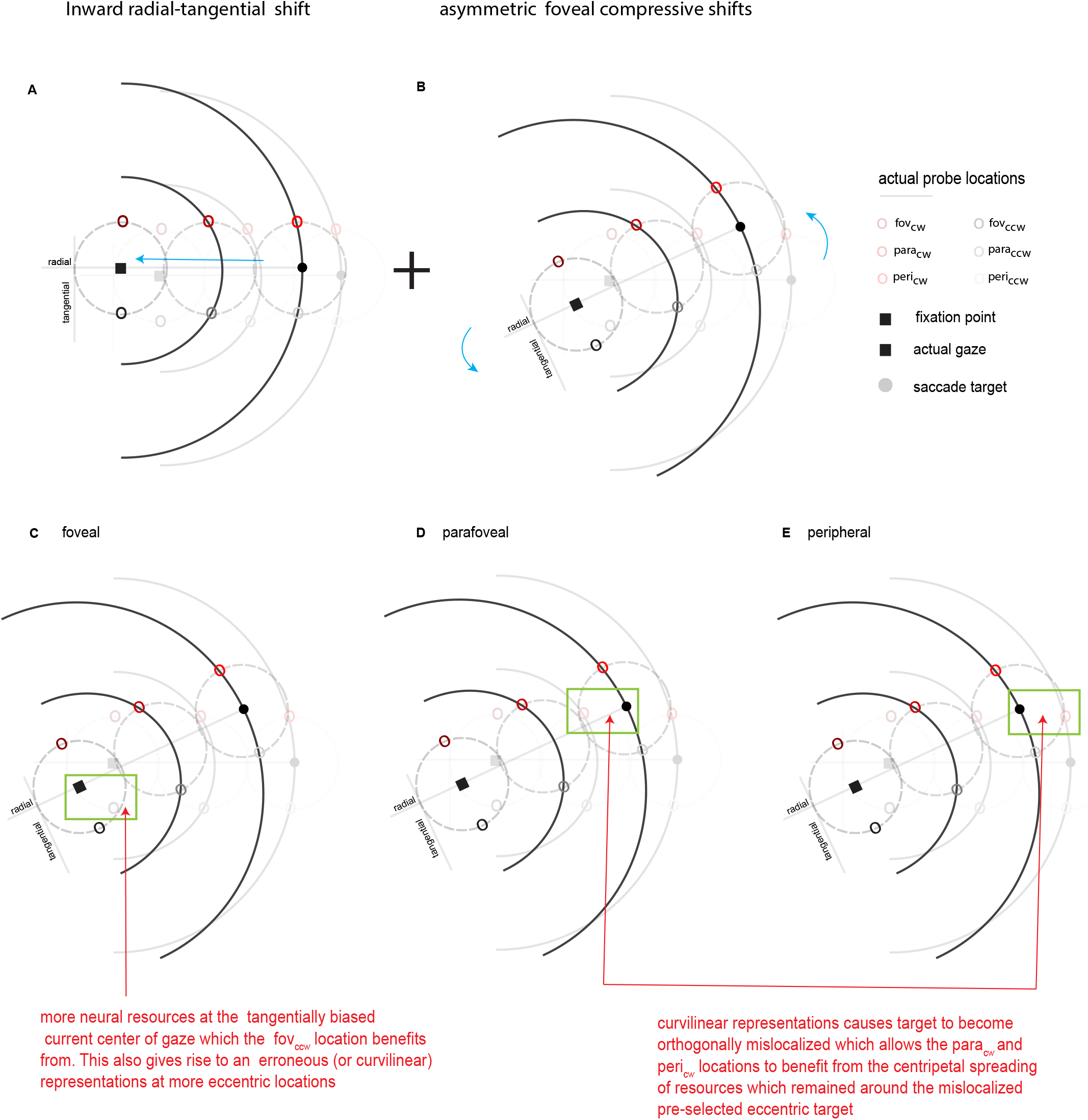
Asymmetric foveal compression. **(A-B)** simple schematic showing the combination of an inward radial-tangential shift towards the fovea and sustained asymmetric foveal compressive shifts. **(C-E)** Visual sensitivity predictions at geometrically symmetric tangential retinotopic locations in the foveal, parafoveal, and peripheral regions of visual space.

### Functional predictions and presaccadic visual sensitivity results

From here on out, we will refer to perimicrosaccadic compression of visual space, which we posit is actively modulated by persistent and sustained asymmetric compressive components as simply “curved visual space”. Curved visual space accompanies very specific and precise presaccadic visual sensitivity predictions at the foveal and more eccentric regions of visual space. Specifically, in the fovea, transient but marked increases in presaccadic visual sensitivity at retinotopic foveal locations that either overlap with or are more proximal to the foveal region that constitutes the target of these persistent and sustained asymmetric compressive shifts (*fov*_*ccw*_ location) should be observed. Furthermore, probes flashed at retinotopic foveal locations (*fov*_*cw*_) that are more distal to the targeted region of these asymmetric compressive shifts should experience marked declines in presaccadic visual sensitivity (Fig. 3C). Conversely, in the more eccentric regions of visual space, we should observe transient but marked increases at eccentric locations that are more proximal to the centripetal spreading of resources that remain around the mislocalized preselected target (*para*_*cw*_ and *peri*_*cw*_, *9, 11*), while eccentric locations (*para*_*ccw*_ and *peri*_*ccw*_) that are more distal should experience marked declines in presaccadic visual sensitivity (Fig. 3D-E).

To directly assess these predictions from a functional perspective, we calculated visual sensitivity functions by binning percent correct responses across subjects for the foveal, parafoveal, and peripheral experiments using a 15 ms sliding window that was moved in 45 ms increments. Next, we obtained each subject’s raw visual sensitivity using a 20-fold jackknife procedure by estimating sensitivity from 95% of the data. This was repeated 20 times, each time leaving out 5% of the data. To then control for any eccentricity-dependent effects across visual space, these traces were normalized by the average raw sensitivity over the initial 100 ms in the data. In the foveal retinotopic region, during persistent and sustained compression toward the *fov*_*ccw*_ retinotopic location, we observed transient but marked increases in presaccadic visual sensitivity at the *fov*_*ccw*_ retinotopic location. At the *fov*_*cw*_ retinotopic location, we observed transient but marked declines in visual sensitivity (Fig. 4A). Indeed, as we had expected, these foveal results are predicted by curved visual space. In the parafoveal region, we observed transient but marked increases in presaccadic visual sensitivity at the *para*_*cw*_ retinotopic location, while at the *para*_*ccw*_ retinotopic location, we observed marked declines in visual sensitivity (Fig. 4B). Finally, in the peripheral region, we observed transient but marked increases in visual sensitivity at the *peri*_*cw*_ retinotopic location, while at the *peri*_*ccw*_ retinotopic location, we observed transient but marked declines in visual sensitivity (Fig. 4C). Similar to the foveal region, these results are also predicted by curved visual space.

**Fig 4.**
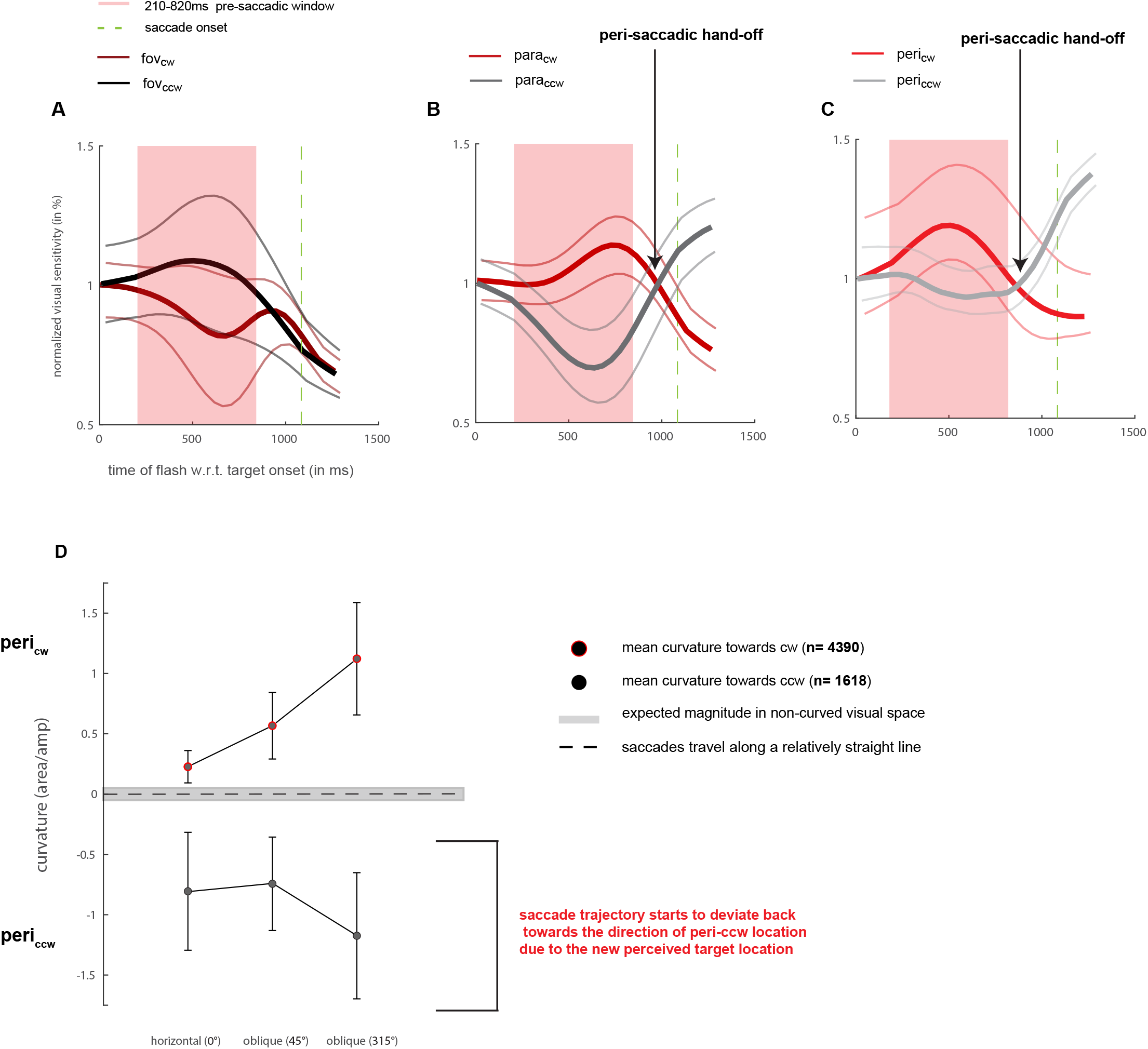
visual sensitivity, and trajectory deviation estimates. **(A-C)** Changes in visual sensitivity along tangential axes with respect to target onset; error estimates were calculated using a 20-fold jackknife procedure. **(D)** Mean saccade trajectory deviation for horizontal and oblique saccades across experiments. Positive values on the ordinate denote trajectory deviation towards a clockwise location (cw), while negative values denote trajectory deviation towards a counterclockwise location (ccw).

### Perisaccadic sensitivity hand-offs in the more eccentric regions

Remarkably, around a perisaccadic window (∼100 ms before saccade onset), we also observed perisaccadic visual sensitivity “hand-offs” between clockwise and counterclockwise locations at the more eccentric regions of visual space. Specifically, 1014 ms after the onset of the fixed eccentric target (i.e., 70 ms before saccade onset), which is around the time human subjects orthogonally mislocalize flashed faint stimuli away from the actual location of the saccade target (*11,12*), the *para*_*ccw*_ retinotopic location that once experienced transient declines in perisaccadic visual sensitivity began to experience rapid increases in perisaccadic visual sensitivity. Conversely, the *para*_*cw*_ retinotopic location that once experienced transient increases in perisaccadic visual sensitivity began to experience rapid declines in perisaccadic visual sensitivity (Fig. 4B). Along the same lines, in the peripheral region, we observed earlier perisaccadic hand-offs between *peri*_*ccw*_ and *peri*_*cw*_ retinotopic locations. Specifically, 920 ms after the onset of the fixed saccade target (i.e., 164 ms to saccade onset), the *peri*_*ccw*_ retinotopic location that once experienced transient declines in perisaccadic visual sensitivity began to experience rapid increases in perisaccadic visual sensitivity, while the *peri*_*cw*_ retinotopic location that once experienced transient increases in perisaccadic visual sensitivity began to experience rapid declines in perisaccadic visual sensitivity (Fig. 4C). As expected, in the foveal retinotopic region, from 820 ms onward after the onset of the fixed eccentric target, we observed, for the most part, global declines in perisaccadic visual sensitivity at both foveal retinotopic locations (positioned away from the future center of gaze, Fig. 4A). Overall, these results suggest that just before the eye was set to move, resources that were asymmetrically compressed in the fovea were slowly reallocated back to more eccentric locations, which allowed these regions to calibrate their representations back toward counterclockwise locations. As we might expect, this calibration (or hand-off) appears to benefit the *para*_*ccw*_ and *peri*_*ccw*_ locations.

### Curved saccade trajectories

Having provided the functional correlates of curved visual space, a natural question one might ask is whether the planning of a saccade, its eventual execution, and the transient changes in visual sensitivity we highlighted share an underlying cause. To investigate this question, we calculated trajectory estimates associated with the saccades subjects made toward the periphery. We predicted that saccades on average would deviate along a curved path toward the *peri*_*cw*_ retinotopic location – the retinotopic location toward which subjects on average mislocalized the fixed saccade target (Fig. 2C). Keeping in mind that the perisaccadic handoffs that occurred to some extent temporally overlapped with when subjects began to plan and prepare for an impending saccade (i.e., ∼150 to 200 ms before saccade onset) (*21 22)*, we further predicted that a smaller number of saccades should also deviate back toward the *peri*_*ccw*_ retinotopic location.

Aligned with both of our predictions, we observed that trajectories toward the periphery systemically deviated along a curvilinear path across experiments, at a rate of 73% toward the *peri*_*cw*_ retinotopic location. In 27% of all trials, saccades were curved toward the *peri*_*ccw*_ retinotopic location. These observations align with previous studies that report that the spatial extent human subjects predict that the saccade target is located actively influences the planned saccade trajectory (*14*). It also aligns with the finding that when a planned saccade is directed toward the incorrect location of the target (e.g., an orthogonal position in space), human subjects are indeed able to calibrate the trajectory of the saccade toward the correct location of the target (*27*). Specifically, when the eccentric target was presented at a fixed azimuth angle of 0°, the trajectories were curved toward the *peri*_*ccw*_ retinotopic location at a rate of 91% with an average curvature of 0.22 in area/amp. Conversely, for the remaining trials (saccades curved toward the *peri*_*ccw*_ retinotopic location), an average curvature of -0.80 in area/amp was observed. When the saccade target was presented at a fixed azimuth angle of 45°, the trajectories were curved toward the *peri*_*cw*_ retinotopic location at a rate of 65% with an average curvature of 0.56 in area/amp, while for the remaining trials (saccades curved toward the *peri*_*ccw*_ retinotopic location), an average curvature of -0.74 in area/amp was observed. Finally, when the saccade target was presented at a fixed azimuth angle of 315°, trajectories were curved toward the retinotopic *peri*_*cw*_ retinotopic location at a rate of 60% with an average curvature of 1.12 in area/amp, while for the remaining trials (saccades curved toward the retinotopic location *peri*_*ccw*_ location), an average curvature of -1.17 in area/amp was observed.

### Rhythmicity supporting local transient shifts in gaze

We next investigated whether visual sensitivity changes in the foveal region compared to the more eccentric regions of visual space were uniquely supported by rhythmicity (∼3.3 Hz), which actively mediate neural signals within attention modulating retinotopic brain areas that predict fixational eye movements (*28*). To do this, we applied the same exact technique used in a seminal study where perceptual data within a 300–1100 ms window were used to explore the relationship between attentional phenomena and global oscillating effects (*29*). However, because we decided to compute our visual sensitivity functions within a 15 ms window shifted by 45 ms increments as opposed to the 15 ms increments used in the foremost study mentioned, we compensated for this difference by using our entire data set, which included presaccadic and perisaccadic sensitivity signatures (0-1285 ms). Consequently, this increased the reliability of our spectral results. Additionally, to ensure that our spectral results revealed the true periodic nature of our data, we detrended our nonnormalized time-series data (Supplementary. Fig. 1). Finally, we applied a Hanning window to control for any spectral leakage and then transformed our detrended data into the frequency domain using a fast Fourier transform (FFT).

At the *fov*_*ccw*_ retinotopic location, we observed that these transient changes in visual sensitivity were actively mediated by rhythmicity in the delta frequency band at ∼2 Hz. This rhythmicity is known to suspend neural resources during cognitively demanding tasks (Fig. 5A, 30). At the *fov*_*cw*_ retinotopic location, we observed that these transient changes in visual sensitivity were actively mediated by rhythmicity at ∼3 Hz, which previous studies have shown within attention modulating retinotopic brain areas predicts the occurrence of fixational eye movements (*29*). Additionally, this rhythmic range is also known to mediate the allocation of attentional signals that support visual sensitivity at suprathreshold levels (31*-33*). Furthermore, the spectral signature that mediated the *fov*_*ccw*_ retinotopic location was not location-specific when compared to shuffled data sets (Fig. 5B, p=0.2280). Conversely, the spectral signature that supported the *fov*_*cw*_ retinotopic location was specific to that retinotopic location (p=0.0230). The *para*_*cw*_, *para*_*cw*_, *peri*_*ccw*_, and *peri*_*ccw*_ retinotopic locations also exhibited rhythmicity in the ∼2 Hz delta frequency band (Fig. 5C & 5E). The spectral signatures observed at these eccentric clockwise and counterclockwise retinotopic locations were not location-specific (p=0.0915, p=0.9725, p=0.3179, p=0.9725) (Fig. 5D & 5F).

**Fig 5.**
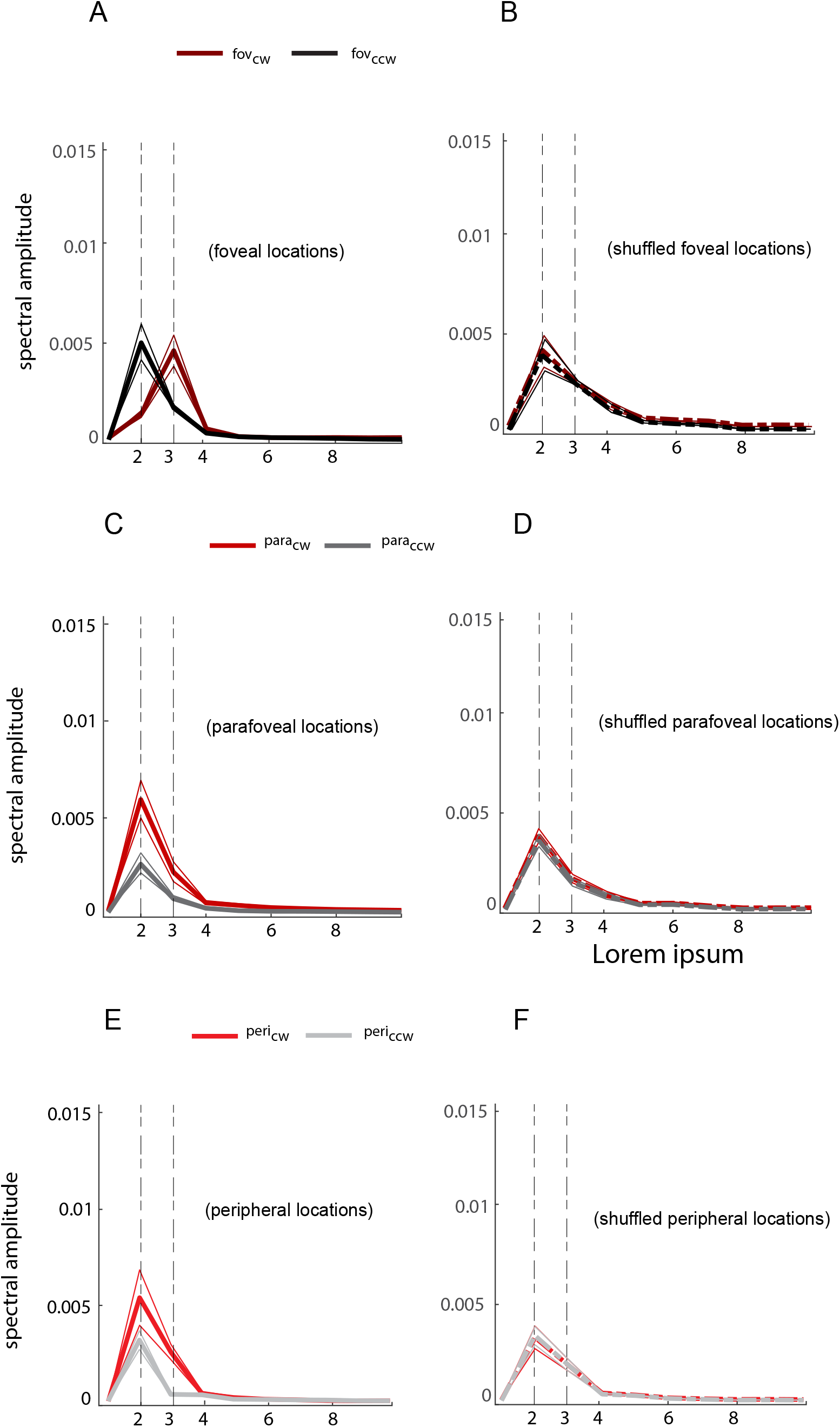
Frequency domain estimates. **(A-F)** Frequency domain representation of raw detrended peri-saccadic sensitivity at less and more eccentric locations in visual space. Shuffled (n=1000, dashed line) versus unshuffled (solid line) data set were used to compute location-specific spectral signatures at a p<0.05.

While the link between attentional signals originating from attention modulating retinotopic brain areas (e.g., the superior colliculus, frontal, parietal, visual cortices) and global oscillating effects in general is debated, empirical evidence is starting to emerge in support of this relationship (28, *30,34,35,36*). Here, our spectral results from a functional perspective further support this emerging idea (*29*). Specifically, our results suggest that while visual resources that would otherwise facilitate active visual processing at the more eccentric retinotopic regions of visual space appear inhibited, marked by delta oscillations (Fig. 5, *30, 37*), visual resources remained present in the fovea specifically around the *fov*_*cw*_ retinotopic location. It is important to note that this retinotopic location was not only proximal to the site where initial transient shifts in gaze were directed toward (*18, 19*) but also constitutes the retinotopic location tasked with transferring its resources toward a foveal region that will become more proximal but not directly aligned to the *fov*_*ccw*_ retinotopic location (Fig. 2B). Despite these results, it is difficult to directly assess the extent to which these global oscillating effects reflect the neural dynamics across attention modulating retinotopic brain areas. To this end, we proposed a phenomenological force field model to investigate whether our model could recapitulate visual sensitivity changes observed at the less eccentric (foveal) and more eccentric (parafoveal) regions of visual space.

### A novel phenomenological force field model

Our model includes two main layers – a “retinotopic field” and a “force field”, both of which are neurobiologically inspired. Our retinotopic field is composed of eccentric configured overlapping retinotopic population receptive fields (pRFs) that tile foveal, parafoveal and peripheral retinotopic regions of the visual space, analogous to the small receptive fields that support foveal representation in the primate visual cortex and increase in size with stimulus eccentricity (Fig. 6A). Additionally, pRFs in our model were assigned population elastic fields (pEFs), which, like our pRFs, were also proportional to their eccentricity. Considering that pRFs across retinotopic brain areas can predictively shift beyond the spatial extent of their classical receptive fields (*38, 39, 40*), pEFs allow our modeled pRFs the ability to undergo transient perturbation beyond their classical surround structures. At the same time, it ensures that pRFs do not go beyond this movement extent, thus allowing the visual system to maintain retinotopic organization.

**Fig 6.**
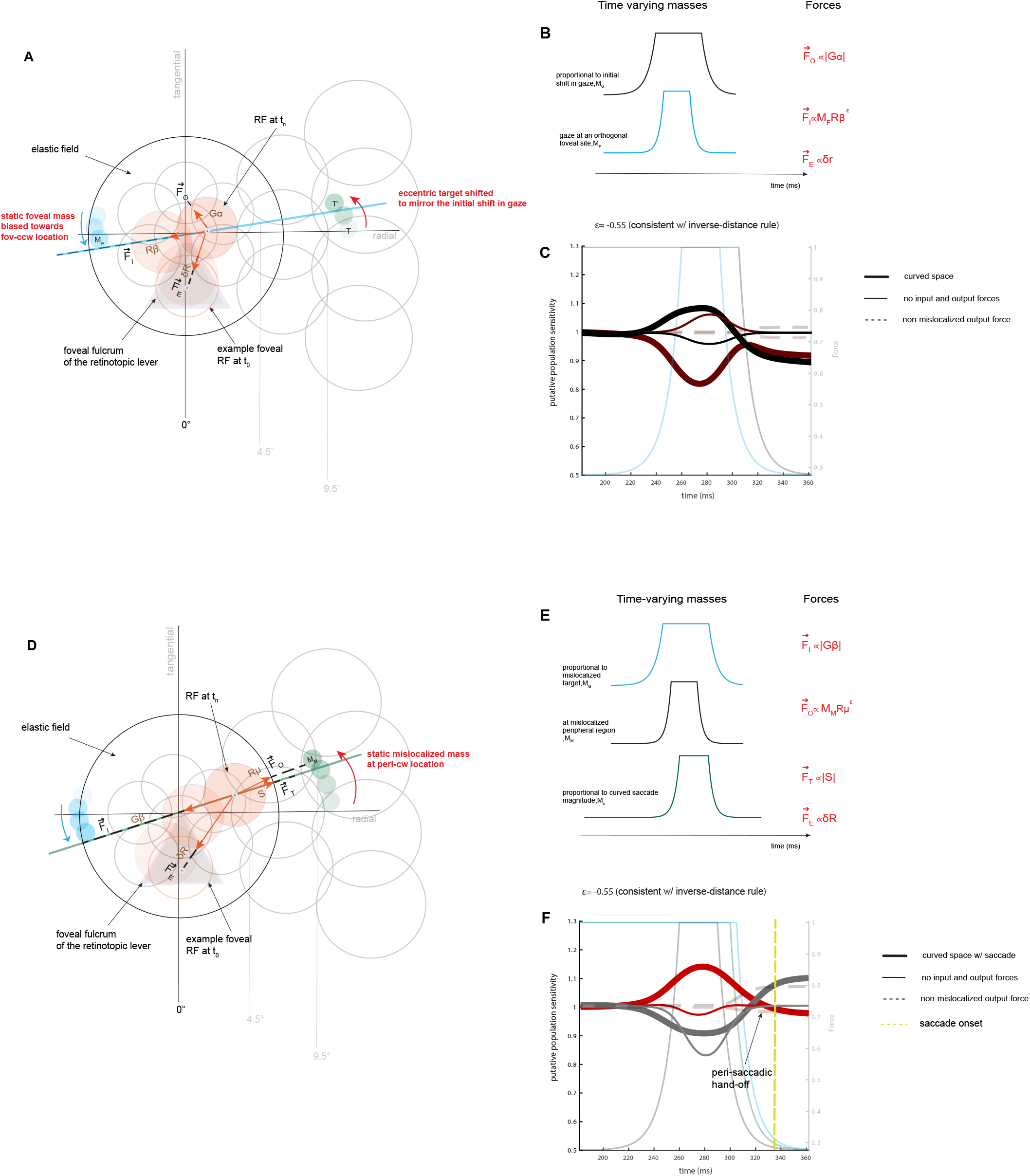
Phenomenological model of curved visual space. **(A)** For illustration purposes, a single eccentricity dependent population receptive field (pRF) within a retinotopic field is highlighted in pink. A pRF is defined by its center, size, and an eccentric dependent population elastic field (pEF). **(B)** M_Gα_ denotes gaze transitional signals modelled as a virtual mass at infinity located in a direction approximating the initial shift in gaze. M_F_ denotes compressive signals modelled as a varying mass directed towards a counterclockwise tangential region in the fovea. **(C)** Simulation results under three main conditions **(D)** A pRF bounded by pEF is highlighted in pink for parafoveal simulations. **(E)** M_Gβ_ denotes gaze translational signals modelled as a virtual mass at infinity located in a direction approximating compressed shift in gaze, M_M_ denotes a varying mass at a mislocalized peripheral region, and M_S_ denotes the curved transitional signals modelled as a virtual mass at infinity located in the direction of a curvilinear saccade. **(F)** simulation results under three main conditions.

The second and final layer of our model is a “force field”. Our force field is founded on three key assumptions, two of which have been well established. The first assumption is that attention mediating retinotopic brain areas during fixation makes available their neural resources, which are used by the visual system to facilitate visual processing at an attended foveal retinotopic site. Furthermore, as the eye is set to move, the same resources these areas provide are reallocated toward more eccentric regions in visual space (*7, 23, 24, 41-48*). The second assumption is that around the time of a saccade, oculomotor signals, which also accompany their own set of resources, originating from premotor brain areas become available. These resources are used by the visual system to enhance the future fields of retinotopic cells (*49-51*).

The third assumption is that attentional-oculomotor signals described above are actively modulated by “transient eccentric error signals”. These signals likely originate within attention modulating retinotopic brain areas that specifically regulate the allocation of resources away and toward the more eccentric regions of visual space (e.g., visual cortices and parietal cortices, *9, 20, 26, 38, 52, 53, 54*). We propose that these signals directly support persistent and sustained asymmetric retinotopic compressive shifts. Together, the summation of attentional-oculomotor signals and transient eccentric error signals give rise to temporally overlapping predictive input and output forces that are exerted by corresponding time varying masses – M. These forces perturb pRFs akin to a mechanical lever or what we refer to as a “retinotopic lever”, thereby distorting visual representation along retinotopic displacements that include an angular bearing. Furthermore, keeping in mind how frequently we saccade (∼ 3-5 times per second), which is energetically expensive (*15*), the final assumption of our force field is that the neural computation that balances energy costs and the distribution of resources within different regions of visual space acts within an inverse force field (*55*). Specifically, this ensured that pRFs were appropriately sensitized toward task-relevant retinotopic locations while minimizing the energy cost associated with spatial perturbations.

### The force field model can explain asymmetric differences in visual sensitivity

Simulations under an inverse force field (*ε*=-0.55), sampled at discrete increments of 1 ms, produced spatiotemporal retinotopic displacements of pRFs (Supplementary. Fig. 2-3), with each pRF modeled as a bivariate Gaussian kernel function (see Methods). With this, we calculated time-varying modulation density read-outs (or putative population sensitivity), equivalent to the visual sensitivity signatures in our psychophysical experiments.

As far as the foveal simulations, modeled pRFs were initially perturbed by the introduction of a radial-tangential translational force (i.e., an output force) exerted by a virtual mass at infinity positioned in the direction of an initial shift in gaze toward a foveal clockwise region (Gα). Under the influence of this output force, to maintain retinotopic organization, pRFs that tile the more eccentric retinotopic regions of visual space were restricted from over-translating toward the fovea. Because of this, the more eccentric pRFs, for the time being, remained slightly displaced (or orthogonally mislocalized) toward a radial-tangential retinotopic site at the more eccentric regions of visual space. Shortly after, we introduced a predictive asymmetric compressive force (i.e., an input force) exerted by a mass positioned in the fovea (Rβ). In addition to radial-tangential translational retinotopic shifts, this caused pRFs to start undergoing transient asymmetric retinotopic compressive shifts toward a foveal counterclockwise region. Under the influence of these temporally overlapping input and output forces, we observed putative population sensitivity signatures that recapitulated changes in visual sensitivity obtained from our psychophysical experiment in the fovea (Fig. 6C, Fig. 4A). Assuming that pRFs were not mediated by transient eccentric error signals (i.e., simulations that did not include an input or output force) and were simply perturbed by oculomotor signals originating from premotor brain areas (i.e., a purely predictive translational force), we observed, as one might have expected, no difference in putative population sensitivity between modeled *fov*_*cw*_ and *fov*_*ccw*_ retinotopic locations (Fig. 6C, dotted marker, *38*). Similarly, retinotopic shifts that only included a non-mislocalized output force (Fig. 6C, thin markers, *7*) were also unable to explain the observed changes in visual sensitivity in the fovea.

Regarding the simulations for the more eccentric parafoveal region (Fig. 6D), an additional predictive asymmetric compressive force (i.e., an input force) was exerted by a mass positioned directly at the center of a foveal counterclockwise region (Gβ). The onset of this input force ensured that pRFs, in a very persistent and sustained manner, remained under the influence of the fovea. Next, we introduced a predictive mislocalized convergent force (i.e., an output force) exerted by a mass positioned around a clockwise region in the periphery (Rμ). Keeping in mind that these forces obey an inverse distance rule, which requires that pRFs further from a given mass give less of its resources than pRFs closer to the same mass, the introduction of this output force ensured that while resources remained sustained in the fovea, some were also centripetally distributed around an eccentric orthogonal site. Overlapping with the onset of this output force was a predictive translational force (analogous to when subjects began to plan for a saccade). This translational force was exerted by a mass positioned in the direction of the impending curvilinear saccade that slowly deviated back toward a counterclockwise region in the periphery (Fig 4C). Note that we modeled the offset of this curvilinear translational force as the onset of a saccade, which in the context of our psychophysical experiments is when the subject stopped planning for the impending saccade and eventually executed a rapid eye movement toward a mislocalized eccentric region. Under the influence of these temporally overlapping forces, we observed presaccadic signatures that quite clearly recapitulated changes in visual sensitivity obtained from our psychophysical experiment in the parafoveal region of visual space (Fig. 6F; Fig. 4B). In addition, we observed a perisaccadic hand-off just before saccade onset. In all, as expected, simulations that did not include the combination of an input and an output force or those that only included a correctly localized output force could not explain the visual sensitivity changes we observed in our psychophysical experiments.

## DICUSSION

Peri-microsaccadic compression of visual space — the ability of retinotopic cells to transiently exhibit predictive spatiotemporal retinotopic compressive shifts toward the target of the impending fixational eye movement — is known to dramatically alter visual perception. However, it was unknown whether perimicrosaccadic compressive shifts, under certain conditions, can become spatially asymmetric, that is, demonstrate a persistent and sustained bias toward one foveal locus over another. If this type of transient top-down modulating occurs, what perceptual and oculomotor effects does it enable?

To provide an account from a functional perspective, we systematically measured the consequences of transient shifts in gaze on visual sensitivity at geometrically symmetric retinotopic locations in the foveal, parafoveal, and peripheral regions of visual space within presaccadic temporal windows. We also assessed how these transient shifts in gaze mediate the planning and execution of a saccade trajectory. We show for the first time that these transient shifts in gaze give rise to local asymmetric differences in visual sensitivity at geometrically symmetric retinotopic locations within a presaccadic window, while saccade trajectories deviate along a curvilinear path toward an orthogonal mislocalized peripheral site (Fig. 4). We later constructed a phenomenological force field model in which we proposed that transient shifts in gaze are underlined by attentional and oculomotor signals that are actively modulated by transient eccentric error signals. The summation of these signals manifests as temporally overlapping predictive input and output forces that act on cells across retinotopic brain areas and thereby modulate their population sensitivity equivalent to changes in visual sensitivity in our psychophysical experiments. Putative population sensitivity signatures recapitulated local asymmetric differences in visual sensitivity we observed at geometrically symmetric retinotopic locations in increasingly eccentric retinotopic regions of visual space (Fig. 6).

Apart from uncovering the effects of transient local shifts in gaze on presaccadic visual sensitivity and saccade trajectory, our behavioral and simulation results provide two fundamental insights into how asymmetric perimicrosaccadic compressive shifts actively modulate perisaccadic compressive representation of visual space. First, perisaccadic representation reflects the influence of attentional-oculomotor signals that are actively modulated by transient eccentric error signals and is not solely the result of purely translational or symmetrically convergent signals, as suggested by previous studies (*7, 38*). Indeed, the presence of or the gating off of these eccentric error signals (modeled as the onset and offset of curvilinear translational signals) seems to determine (i) whether flashed stimuli will be orthogonally mislocalized away from or toward a fixed saccade target (*8, 11, 12*), (ii.) and whether retinotopic cells will align with or shift along an axis that is orthogonal to the radial axis (*9,10*). Finally, the neural computations that support this retinotopic representation well before and around the time of a saccade obey an inverse-distance rule, while the neural architecture that mediates spatiotemporal retinotopic shifts pRFs undergo – which give rise to this representation – includes pEFs. Indeed, these critical insights should inform future neural investigations that explore the relationship between presaccadic and perisaccadic representations of visual space.

To conclude, the observed transient local shifts in gaze during fixation remain a curious point since they are directed away from the fixed eccentric target. However, considering that humans were once prey, it is possible that these miniature shifts away from the visual target of interest reflect an evolutionarily conserved visual oculomotor avoidance or escape strategy the mammalian visual system used to track a distal target (e.g., conspecific, predator) from a safe distance (e.g., corner of the eye), later followed by a calculated movement (e.g., saccade) toward the retinotopic location previously avoided (*56-58*).

## METHODS

### Human subjects

The protocol for this study and the collection and storage of the data were approved by the Yale Ethics Review board and were in accordance with the Declaration of Helsinki. All subjects gave informed constent. Subjects all above 18 years of age gave both verbal and written consent. Specifically, six consenting human subjects (5 females and 1 male; 21-26 years old) with corrected-to-normal visual acuity participated in the three experiments reported as it is standard within the field of research. Three of the six subjects participated in the foveal and parafoveal experiments. The remaining three subjects participated in the peripheral experiment. The Yale Ethics Review board approved the protocol of these experiments and the storage of the data that were collected.

### Visual stimuli and experimental protocol

Fig. 1 summarizes the protocol used across experiments. In a dimly lit room, stimuli were presented on a gamma corrected display with a spatial resolution of 1400 × 1050 pixels and a mean luminance of 38 cd/m2. Eye movements were recorded using an infrared video-based eye tracker sampled at 1 kHz (I-Scan Inc., Woburn, MA). While tightly head fixed using a head rest and a customized bite bar, subjects at the onset of each trial acquired fixation for 300 ms. Fixation within this temporal window was spatially aligned with the fixation point (0.5 dva in diameter) using a custom-developed 9-point eye-calibration procedure in Picto. Next, depending on the subject, a saccade target was presented at an eccentricity of 10° and at azimuth angles of 0° (for *s*_1_), 45° (for *s*_2_) and 315° (for *s*_3_) (Fig. 1A). Following target onset, fixation was maintained for another 300-600 ms within an ∼2 dva fixation tolerance window, at the end of which the fixation spot was extinguished, which served as the central motor cue for subjects to make an eye movement toward the saccade target. For another 500 ms after the central motor cue, the saccade target remained visible. Finally, a low-contrast probe (at a contrast level that the subject could detect with 50% accuracy) was isotropically flashed, depending on the experiment, for 20 ms randomly either along the radial or tangential axis at a random time during the 300-600 ms window between the target onset and the central motor cue (25% of trials) or at a random time between 0-340 ms after the central motor cue (50% of the trials). In the remaining 25% (which yielded a false alarm rate of 1.2%), no probes were presented. All three conditions were randomly interleaved across trials. With respect to target onset, the presentation of flashed probes fell within a time window of 0-1285 ms after target onset (Fig. 1B). Subjects were asked to report, using a push button, whether they were able to detect the flashed probe. While the responses along the radial axis were not analyzed, the presentation of these probes was crucial in controlling for any response biases that may have emerged along the tangential axes. To ensure that attentional and oculomotor resources were not driven by an asymmetric stimuli configuration, as is the case in previous studies, no secondary salient spatial cue was presented. Across experiments, subjects, and axes, we collected a total of ∼ 18,000 trials, while along the tangential axes, a total of ∼6010 trials were collected.

### Statistical analyses

To investigate gaze dynamics, probability density maps and variance estimates were calculated using median filtered edge preserving fixational data, which were rotated to an angle of 0° across target locations. Next, we calculated a gaze probability density map for the foveal and peripheral regions using a bivariate Gaussian kernel (*59*). The estimated locus of gaze across subjects was defined as the peak of each probability density map (i.e., the yellow contour lines). To compare the distribution of gaze in the fovea during the 0- to 209-ms window after target onset versus the 210- to 820-ms window after target onset, a Mann–Whitney U statistic was calculated at a p value less than 0.05. Furthermore, variance in gaze was calculated using a bivariate contour ellipse area (BCEA) approach that included 68% of fixational data around the mean (*63*). Accompanying error bars were obtained, which denote the standard error of the mean variance across subjects. To investigate the extent, saccade trajectories were curved across experiments at each saccade location. Horizontal and vertical eye positions, with the time window starting point constituting the moment fixation was maintained to the moment the eye landed in the periphery, were isolated. Next, to obtain a measurement of curvature, the curved area that emerged from the eye position during this time window was calculated. To then obtain a ratio measure of curvature per unit amplitude, this area was divided by the amplitude of the saccade (*60*). Accompanying calculated error bars denote the standard error of the mean at each target location across experiments. To investigate normalized changes in visual sensitivity across subjects, I first binned each subject’s percent correct responses for the foveal (Exp. 1), parafoveal (Exp. 2), and peripheral (Exp. 3) experiments using a 15 ms sliding window that was moved in 45 ms increments. Next, we obtained each subject’s raw visual sensitivity using a 20-fold jackknife procedure by estimating sensitivity from 95% of the data. This was repeated 20 times, each time leaving out 5% of the data. This produced a mean and an error estimate of visual sensitivity at each tangential location for each experiment across subjects (*61*). To control for any eccentric-dependent effect on raw visual sensitivity, these traces were normalized by the average sensitivity over the initial 100 ms in the data. To investigate frequency domain and cross-location phase coherence dynamics, a fast Fourier transform (FFT) was applied to the detrended raw visual sensitivity data. Next, we computed the mean of the spectral signatures and corresponding standard error of the mean across subjects. To assess whether the observed periodicities were location specific, we randomly shuffled (n=1000) the probe location identities across trials and calculated the sensitivity function and periodicity of the shuffled data using the same procedures noted above. A two-tailed paired-sample t test at p values of 0.05 was calculated to assess the significance between shuffled and non-shuffled data. Finally, for the cross-location phase coherence analysis, we applied an approach to the shuffled and non-shuffled detrended data that is commonly used in assessing the underlying coherence in neural data (*62*). I then projected the difference between phase angles of each subject between counterclockwise (ccw) and clockwise (cw) locations and then calculated the resultant vector across subjects.

### Novel computational framework - Retinotopic mechanics

The underlying architecture of this simple yet powerful mechanical model is composed of a retinotopic field (*ϕ*_*r*_) and a neurobiologically inspired force field (*ϕ*_*f*_). The retinotopic field is a two-dimensional overlapping hexagonal grid of eccentricity-dependent receptive fields RF_i_ that tiles the foveal, parafoveal and peripheral regions in visual space (Supplementary Fig. 2-3). Each RF_i_ was defined by three parameters: (a) *p*_*i*_, the spatial extent of the RF center (*x*_*i*_, *y*_*i*_) at t_0_ (when there is no resultant predictive force acting on the *ϕ*_*r*_). *p*_*i*_ determines the eccentricity - *e*_*i*_ of RF_i_, (b) *s*_*i*_, the radius of RF_i_ is *prop*ortional to *e*_*i*_, and (c) the movement extent (*M*_*max*_) of RF_i_ is its EF_i_, which is also ∝ *e*_*i*_. *ϕ*_*f*_ emerged from attentional and oculomotor signals actively modulated by transient eccentric error signals that act as predictive input and output forces, and some cases a translational force, which transiently perturbs a constellation of population receptive fields from equilibrium and thereby modulates their putative population sensitivity (density readouts) as a function of space and time. For simulations at the less eccentric (“foveal”), the resultant force 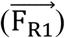 included an input and output force (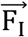, and 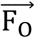) and an internal force 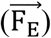, which RF_i_ at t_0+n_ experiences:

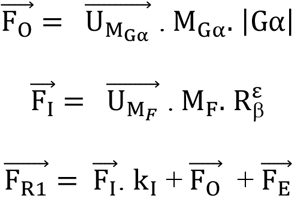

where M_Gα_ and M_F_ are the virtual mass at infinity in the upward direction, which approximates an initial shift in gaze, and a foveal mass located at a lower tangential foveal retinotopic site, which approximates a compressed shift in gaze. 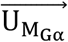 and 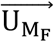 are the unit vectors in the direction of the virtual mass and the foveal mass, respectively. |Gα| is the magnitude in gaze away from the upper tangential foveal retinotopic site, while R_β_ is the spatial difference between RF_i_ and M_F_ raised to an epsilon scalar – ε. The value of ε, depending on the retinotopic mechanics that best explains the phenomenon in question, can either be consistent with a proportional-distance rule (i.e., >0) or an inverse-distance rule (i.e., <0). Furthermore, k_I_ was obtained by calculating the average of 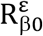 across RF_j_s and then taking its reciprocal 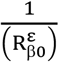, while 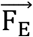 is ∝ the movement of RF_i_. For the more eccentric (“parafoveal”) simulations, the resultant force 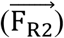 included an input, output, an additional orthogonally biased translational force (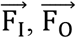 and 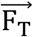) and an internal force 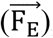, which RF_i_ at t_0+n_ experiences:

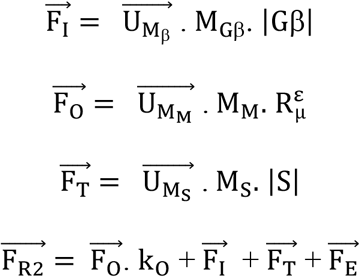

where M_Gβ_ and M_M_ are the virtual mass at infinity at the lower foveal tangential retinotopic site, which approximates a complete compression of gaze, and the mislocalized mass located at a peripheral retinotopic site, which approximates a mislocalized saccade target. 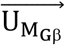 and 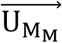 are the unit vectors in the direction of the virtual mass and the mislocalized peripheral mass, respectively. |Gβ| is the magnitude of a fully compressed gaze, which is roughly ∝ the magnitude of a mislocalized target. R_μ_ is the spatial difference between RF_i_ and M_M_ raised to ε, while |S| approximates the magnitude of an impending curvilinear saccade toward the counterclockwise location in the periphery. The same calculation used to obtain k_I_ was used to obtain k_O_; however, in this case, the average of 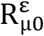 across RF_i_s was used. Depending on the stimulation, a time-varying resultant force (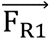 or 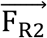) perturbed each RF_i_, which produced spatiotemporal retinotopic displacements (Supplementary Fig. 2-3). The population movements of the constellations of RF_i_s manifested in time-varying modulation of raw density, modeling each RF_i_ as a bivariate Gaussian kernel function *G (64)*. A probability density estimate (or putative population sensitivity readouts), 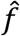, was obtained using the following equation:

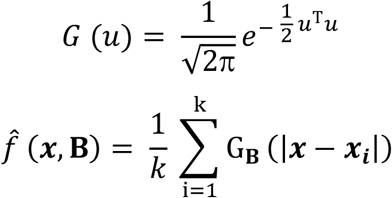

Finally, population sensitivity readouts were later normalized using the same approach used to normalize the raw visual sensitivity traces.

## AUTHOR CONTRUBITIONS

EL conceptualized the project. EL collected the data. EL preprocessed the behavioral data with the assistance of XZ. EL analyzed the behavioral data. EL developed the computational model. EL implemented the model. EL performed the model simulations. EL wrote the manuscript.

## ACKNOWLEDGEMENTS

We would like to thank D. P. Melcher and S. Wirth for comments on the manuscript and B. Wichterlová for editing the manuscript. We would also like to thank A. S. Nandy and M. Scudder for their guidance on the experimental design and implementation.

## DATA AVAILABILITY

The data that support this study are available upon reasonable request.

## CODE AVAILABILITY

The custom MATLAB code that supports the figures reported in this study is available upon reasonable request.

