## Supplementary figures and images for "Visual space curves before eye movements"

### Supp. Fig.1

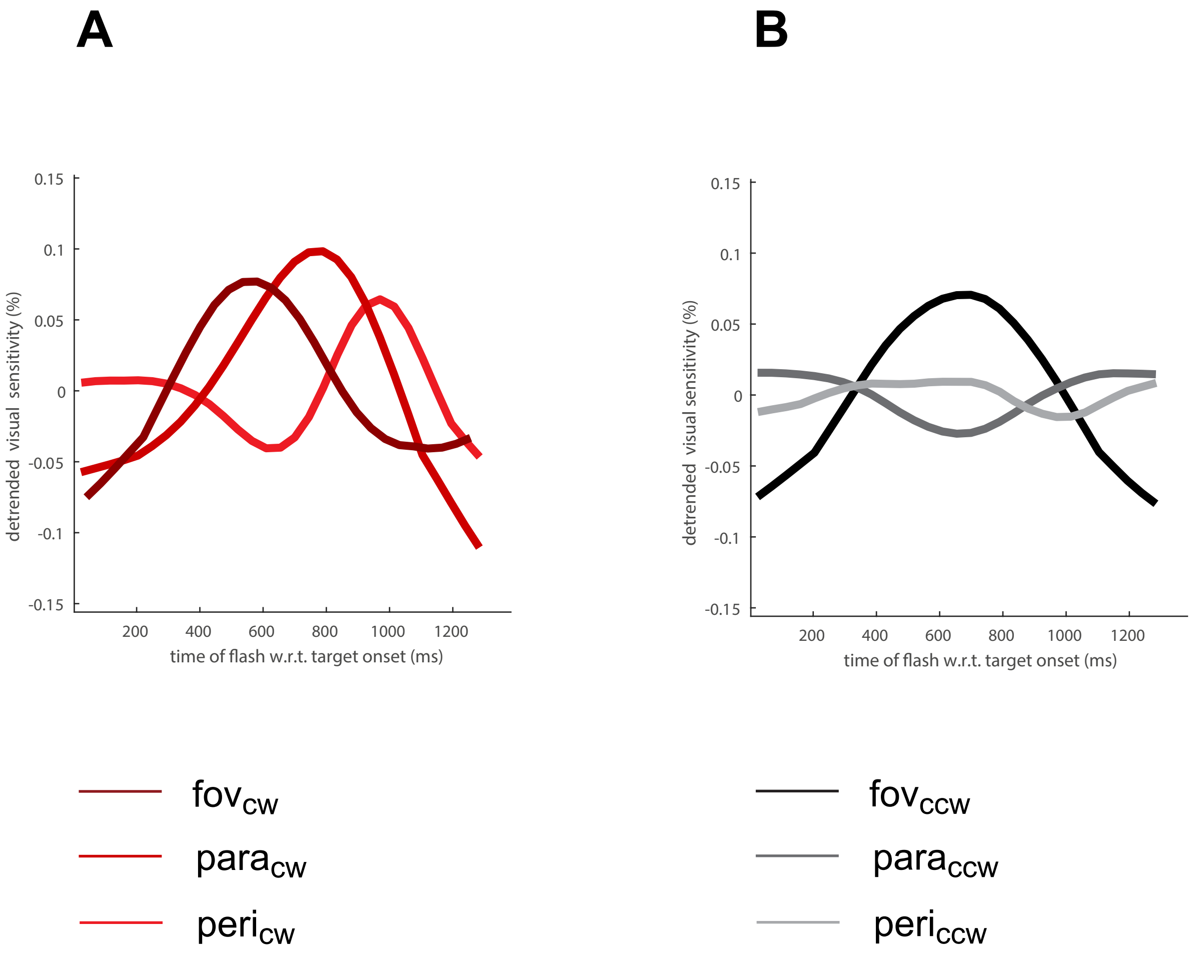

### Supp. Fig.2

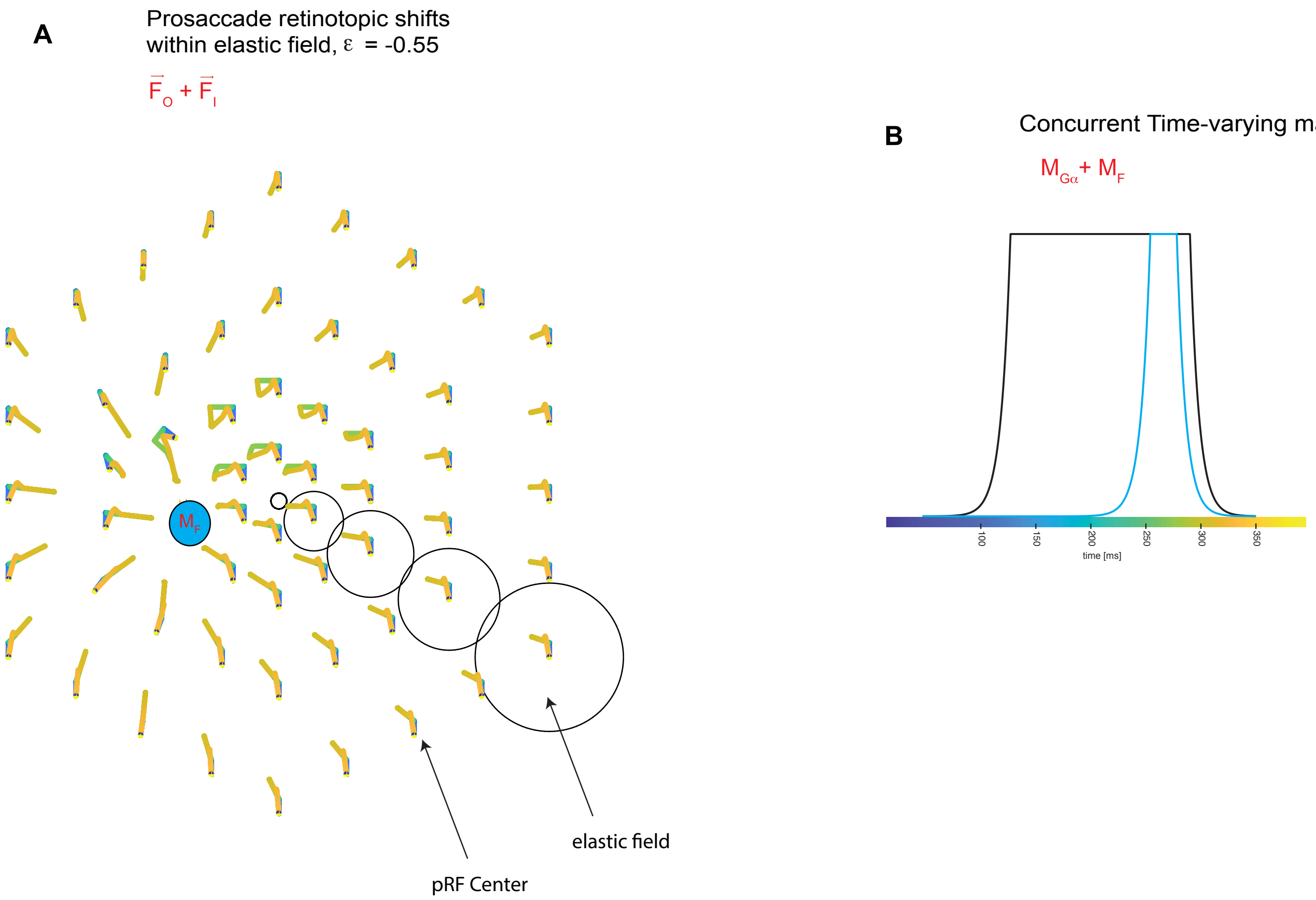

### Supp. Fig.3

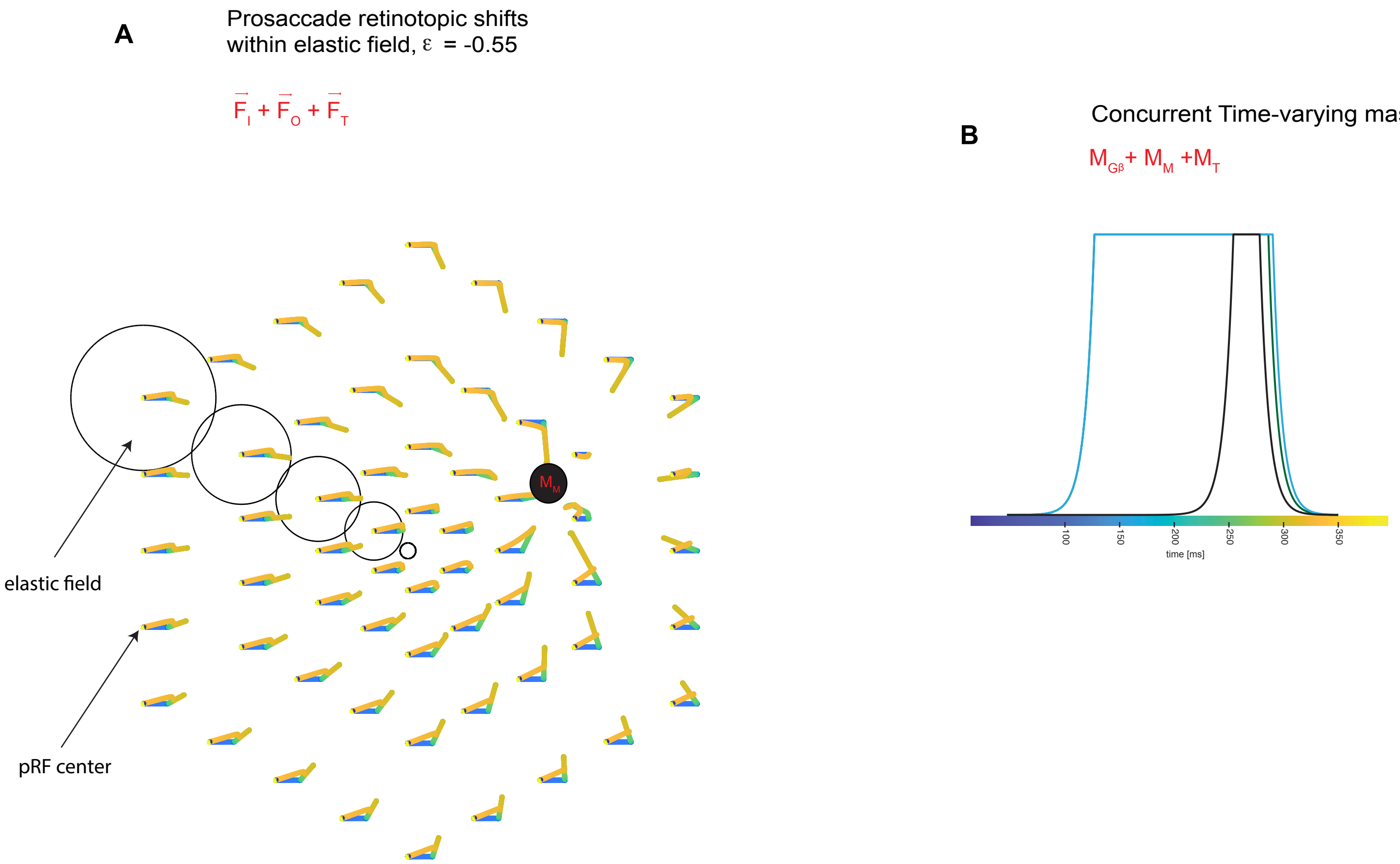
